# Contribution of telacebec to novel drug regimens in a murine tuberculosis model

**DOI:** 10.1101/2024.06.27.601059

**Authors:** Oliver D. Komm, Sandeep Tyagi, Andrew Garcia, Deepak Almeida, Yong Chang, Si-Yang Lee, Jennie Ruelas Castillo, Paul J. Converse, Todd Black, Nader Fotouhi, Eric L. Nuermberger

## Abstract

The clinical efficacy of combination drug regimens containing the first generation diarylquinoline (DARQ) bedaquiline in the treatment of multidrug-resistant tuberculosis has validated ATP synthesis as a vulnerable pathway in *Mycobacterium tuberculosis*. New DARQs in clinical development may be even more effective than bedaquiline, including against emerging bedaquiline-resistant strains. Telacebec (T) is a novel cytochrome bc_1_:aa_3_ oxidase inhibitor that also inhibits ATP synthesis. Based on its demonstrated efficacy as a monotherapy in mice and in a phase 2a clinical trial, we used an established BALB/c mouse model of tuberculosis (TB) to test the contribution of T to novel combination therapies against two strains of *M. tuberculosis* (H37Rv and HN878) in an effort to find more effective regimens. Overall, T was more effective in regimens against the HN878 strain than against the H37Rv strain, a finding that supports the greater vulnerability of the former strain to T and to genetic depletion of QcrB. Against both strains, combinations of a DARQ, clofazimine (CFZ), and T were highly bactericidal. However, only against HN878 did T contribute synergistically, whereas an antagonistic effect was observed against H37Rv. These results demonstrate the therapeutic potential of T and highlight how differences in the susceptibility of *M. tuberculosis* strains could lead to different conclusions about a drug’s potential contribution to novel drug regimens.

## Introduction

Mycobacterial respiration and oxidative phosphorylation have proven to be a vulnerable pathway in *Mycobacterium tuberculosis* for chemotherapy of tuberculosis (TB) (1, 2). Recent drug discovery work has focused on validating potential drug targets, developing small molecule inhibitors and finding combination therapies leading to synergistic effects on the pathway (3-5). Bedaquiline (BDQ, B), a diarylquinoline (DARQ) inhibitor of ATP synthase, was approved for TB treatment by the U.S. Food and Drug Administration in late 2012 (6). Its strong sterilizing activity underpins the efficacy of the first oral 6-month regimen recommended by WHO for treatment of rifampin-resistant TB based on the combination of BDQ, pretomanid (Pa) and linezolid (L), with or without addition of moxifloxacin (M) (regimens abbreviated as BPaL(M)) (7-10). BDQ is a key component of even shorter experimental regimens also containing pyrazinamide (PZA, Z), such as the BPaMZ regimen studied in the recently completed SimpliciTB trial (ClinicalTrials.gov identifier NCT03338621) and the combination of BLZ plus isoniazid and ethambutol studied in the TRUNCATE-TB trial (11). BDQ is also a crucial component of other regimens with strong sterilizing activity in mouse models of TB. For example, BDQ is more effective than rifapentine (P) when either is combined with MZ in BALB/c mice; and the addition of rifabutin (Rb) to BMZ further increases the sterilizing activity (12). The resultant BMZRb regimen will be evaluated in the ongoing phase 2c CRUSH-TB trial (NCT05766267). In addition, the combination of BDQ with the Rv1625c activator GSK2556286 (G) (now in phase 1 of clinical development [NCT04472897]) and the DprE1 inhibitor TBA-7371 (A) (now in phase 2 of clinical development [NCT04176250]), has demonstrated sterilizing activity approaching that of BPaL in BALB/c mice and may represent a promising backbone for further regimen development (13).

Despite its clinical utility, BDQ has several liabilities, including high lipophilicity (e.g., partition coefficient between n-octanol and water [logP] = 7.25 log_10_) and accumulation in tissues with a consequently long terminal half-life of 5-6 months, high plasma protein binding (>99.9%), slow diffusion into caseous lesions, and potential to prolong the QT interval (14-16). Furthermore, although high-level resistance to BDQ caused by mutations in the *atpE* gene have been encountered only rarely in the clinic to date (17), mutations in *Rv0678* (also known as *mmpR5*) (18-20), which cause smaller but still clinically significant reductions in susceptibility are increasingly reported (21-23). One issue of concern is that these mutations are also associated with reduced susceptibility to other TB drugs, such as clofazimine (CFZ) and new DprE1 inhibitors in clinical development (namely TBA-7371 (A) (24), quabodepistat (formerly known as OPC-167832) (25), and the benzothiazinones BTZ043 and PBTZ169 (26)), and these mutations may be observed in isolates from patients without prior BDQ or CFZ exposure (27, 28).

To improve upon the efficacy and liabilities of BDQ, two new DARQ analogues, TBAJ-587 (S587) and TBAJ-876 (S876), with higher potency against *M. tuberculosis* (including *Rv0678* mutants), but lower lipophilicity and improved cardiac safety profiles, were developed (16, 29-31). Mouse efficacy studies have demonstrated their superior potency compared to BDQ against wildtype *M. tuberculosis* H37Rv and isogenic *Rv0678* mutants (18, 32) and superior sterilizing activity when replacing BDQ in combination with Pa and an oxazolidinone (33). Both new DARQs are now in clinical trials (NCT04890535 and NCT06058299).

Telacebec (T) is a new first-in-class anti-mycobacterial drug candidate that binds QcrB to inhibit the cytochrome bc_1_:aa_3_ complex, a terminal oxidase of the electron transport chain (ETC) (34), and therefore acts on a different component of the respiratory chain than BDQ to inhibit ATP synthesis. T shows potent inhibitory activity against *M. tuberculosis* and exceptionally potent bactericidal activity against *M. tuberculosis* mutants (e.g., Δ*cydAB* and *cydC::aph* mutants) lacking the alternative cytochrome bd terminal oxidase and against *Mycobacterium ulcerans* strains which lack a functional *cydAB* operon encoding this oxidase (34-37). Mouse studies testing single doses of up to 1000 mg/kg and human studies testing doses of up to 320 mg/d for 14 days showed favorable safety and tolerability (34, 38-41). A recent Phase 2a study demonstrated dose-dependent early bactericidal activity in TB patients (40). However, there are only limited published data on the potential contributions of T or other QcrB inhibitors to the efficacy of novel drug regimens in animal models of TB chemotherapy to inform regimen development (42).

The fat soluble riminophenazine dye CFZ was originally developed to treat TB in the 1960s but was primarily used as an anti-leprosy medication until recently (43, 44). It was newly designated a second-line drug for rifampin-resistant TB by the WHO, after studies demonstrated the efficacy of 9-month CFZ-containing regimens (45, 46). The mode of action of CFZ is not firmly established, but it has been shown that the primary NADH:quinone oxidoreductase in the mycobacterial ETC, NDH-2, reduces CFZ, which in turn reduces O_2_, recycling CFZ and generating reactive oxygen species (ROS) (47). Lamprecht et al. demonstrated the synergistic activity of BDQ, T and CFZ *in vitro*, finding that this combination sterilized *M. tuberculosis* H37Rv cultures after 5 days of exposure. They hypothesized that the mechanism of the synergy is based on the combined effects of BDQ and T leading to a compensatory increase in electron flux in the ETC and a resulting increase in reactive oxygen species production through CFZ-mediated redox cycling (48).

Taken together, these promising results prompted us to investigate whether T, as an ATP synthesis inhibitor acting on a target different from BDQ, could contribute to novel drug combinations capable of shortening TB treatment and/or mitigating BDQ resistance. Here, we describe a series of experiments to test the hypotheses that T could replace BDQ in, or augment the activity of, DARQ-containing regimens, including combinations of a DARQ, T and CFZ in an established BALB/c mouse model of TB using the H37Rv reference strain (Lineage 4, Euro-American) of *M. tuberculosis*.

The contribution of T to selected combinations was also assessed against an infection with *M. tuberculosis* HN878, a more recent clinical isolate belonging to the other major lineage of *M. tuberculosis* isolates (Lineage 2, East Asian). Compared to H37Rv, HN878 has a lower basal expression of the bd oxidase and is more vulnerable to conditional silencing of *qcrB* expression and T (49). Therefore, we hypothesized T would be more effective in drug regimens against the HN878 strain.

## Results

### Experiment 1: Evaluating the contribution of telacebec when added to core components of, or substituted for bedaquiline in, regimens of clinical significance

Experiment 1 was designed to test (1) whether T has additive activity with drug combinations that have previously been shown to partner well with BDQ, and (2) whether T is as effective as BDQ in these regimens. Therefore, we tested both T and BDQ as additions to each of the following combinations: PaL (50), PaMZ (51), MZRb (12), and GSK2556286+TBA-7371 (13). An additional objective was to determine if the addition of T increases the bactericidal activity of the highly sterilizing BMZ (51, 52) and PMZ (53) combinations.

The experimental scheme is shown in **Table S1**. BALB/c mice received a high-dose aerosol infection with *M. tuberculosis* H37Rv and treatment started two weeks post-infection. Arms containing PZA were limited to 4 weeks of treatment because CFU counts were expected to be very low or undetectable after longer treatment durations and less likely to permit assessment of the contribution of T. Other arms were evaluated after both 4 and 8 weeks of treatment. After 4 weeks of treatment, the addition of T significantly increased the bactericidal activity of PaMZ (p<0.05) and MZRb (p<0.0001) (**Table 1**), but the addition of T did not increase the activity of BMZ although BMZQ was as effective as BMZRb and was superior to BPaMZ (p<0.0001). The addition of T did not increase the activity of PMZ. After 8 weeks of treatment, addition of T antagonized the bactericidal activity of PaL (p<0.01), but did not significantly alter the activity of GA (p=0.78). However, the addition of both T and CFZ to GA significantly reduced the CFU counts compared to GA alone.

**Table 1.** Mean CFU counts at week 4 and week 8 in Experiment 1 (H37Rv)

| | Time point and mean (±SD) $\log_{10}$ CFU counts | | | |
| --- | --- | --- | --- | --- |
| Regimen | W-2 | D0 | W4 | W8 |
| Untreated | 4.61 ± 0.19 | 7.69 ± 0.33 | | |
| Pa<sub>50</sub>L<sub>100</sub> | | | | 4.81 ± 0.46 |
| B<sub>25</sub>PaL | | | | 1.13 ± 0.24 |
| T<sub>5</sub>PaL | | | | 5.86 ± 0.24 |
| PaM<sub>100</sub>Z<sub>150</sub> | | | 6.59 ± 0.22 | |
| BPaMZ | | | 4.51 ± 0.41 | |
| TPaMZ | | | 5.91 ± 0.36 | |
| MZRb<sub>5</sub> | | | 5.75 ± 0.15 | |
| BMZRb | | | 2.78 ± 0.40 | |
| TMZRb | | | 4.53 ± 0.16 | |
| BMZ | | | 2.38 ± 0.05 | |
| BMZT | | | 2.64 ± 0.33 | |
| PMZ | | | 4.97 ± 0.62 | |
| PMZT | | | 4.78 ± 0.13 | |
| 286<sub>35</sub>A<sub>200</sub> | | | | 6.84 ± 0.15 |
| B286A | | | | 1.44 ± 0.17 |
| T286A | | | | 7.14 ± 0.32 |
| TC<sub>6.25</sub>286A | | | | 3.70 ± 0.83 |

Number in subscript indicate dose in mg/kg.
Abbreviations: Pa=pretomanid; L=linezolid; B=bedaquiline; T=telacebec; M=moxifloxacin, Z=pyrazinamide; Rb=rifabutin; 286=GSK-286; A=TBA-7371; C=clofazimine.
W-2 = Day after infection; D0 = Day of treatment initiation; W4/8 = 4/8 weeks after treatment initiation, respectively.

In every example in which either BDQ or T was added to the same 2- or 3-drug combination, BDQ was a stronger contributor to the regimen than T. Addition of BDQ resulted in significantly lower mean CFU counts in all cases, whereas the addition of T only significantly lowered the mean CFU of the MZRb and PaMZ combinations and antagonized the PaL combination.

### Experiment 2: Evaluating the bactericidal activity of regimens based on a backbone of TBAJ-876, clofazimine and telacebec

Experiment 2 was performed to determine if the addition of T to the more potent DARQ TBAJ-876 and CFZ would recapitulate the synergy observed *in vitro* (18, 33). A second objective was to determine if the addition of various fourth drugs would increase the bactericidal activity of this 3-drug combination. Among the fourth drugs tested were the rifamycin transcription inhibitor rifabutin (Rb); three oxazolidinone translation inhibitors: linezolid (L), sutezolid (U), and TBI-223 (O); three cell wall synthesis inhibitors: Pa, the DprE1 inhibitor TBA-7371 (A), and the MmpL3 inhibitor MPL-446 (Mp); and PZA. The experimental scheme is described in **Error! Reference source not found.**. BALB/c mice received a high-dose aerosol infection with *M. tuberculosis* H37Rv and were initiated on treatment 2 weeks post-infection. Treatment duration was limited to 4 weeks because CFU counts in mice treated with TBAJ-876 and CFZ with or without T were expected to be very low or undetectable at 8 weeks.

TBAJ-876 alone at 6.25 mg/kg was highly bactericidal, reducing the CFU counts by more than 3.5 log_10_ over 4 weeks (**Table 2**). Monotherapy with TBAJ-876 was as active as the combination of TBAJ-876 with PaL, as previously observed (18). The addition of CFZ to TBAJ-876 significantly improved the bactericidal activity (p<0.0001). However, addition of T to TBAJ-876+C resulted in significantly higher CFU counts (p<0.01). Driven by the combined additive effects of TBAJ-876 and CFZ, this combination, with or without addition of T, was significantly more active than the TBAJ-876+PaL control regimen (p<0.0001), as was nearly every 4-drug combination based on TBAJ-876 plus CT at this time point. However, only the addition of PZA significantly increased the activity of the TBAJ-876+CT backbone (p<0.001), reducing the CFU counts below the lower limit of detection of 1.66 log_10_ in this experiment. The addition of GSK2556286 to TBAJ-876+CT resulted in the next largest reduction in mean CFU counts, rendering the lungs of two mice culture-negative, but the difference with TBAJ-876+CT was not statistically significant. Neither addition of Rb nor addition of any of the 3 oxazolidinones tested significantly changed the CFU counts after 4 weeks of treatment. With respect to cell wall inhibitors, TBA-7371 was more effective in the combination than the mycolic acid synthesis inhibitors Pa and MPL-446, although only the antagonistic effect of adding Pa to the 3-drug regimen was statistically significant (p<0.0001).

**Table 2.**
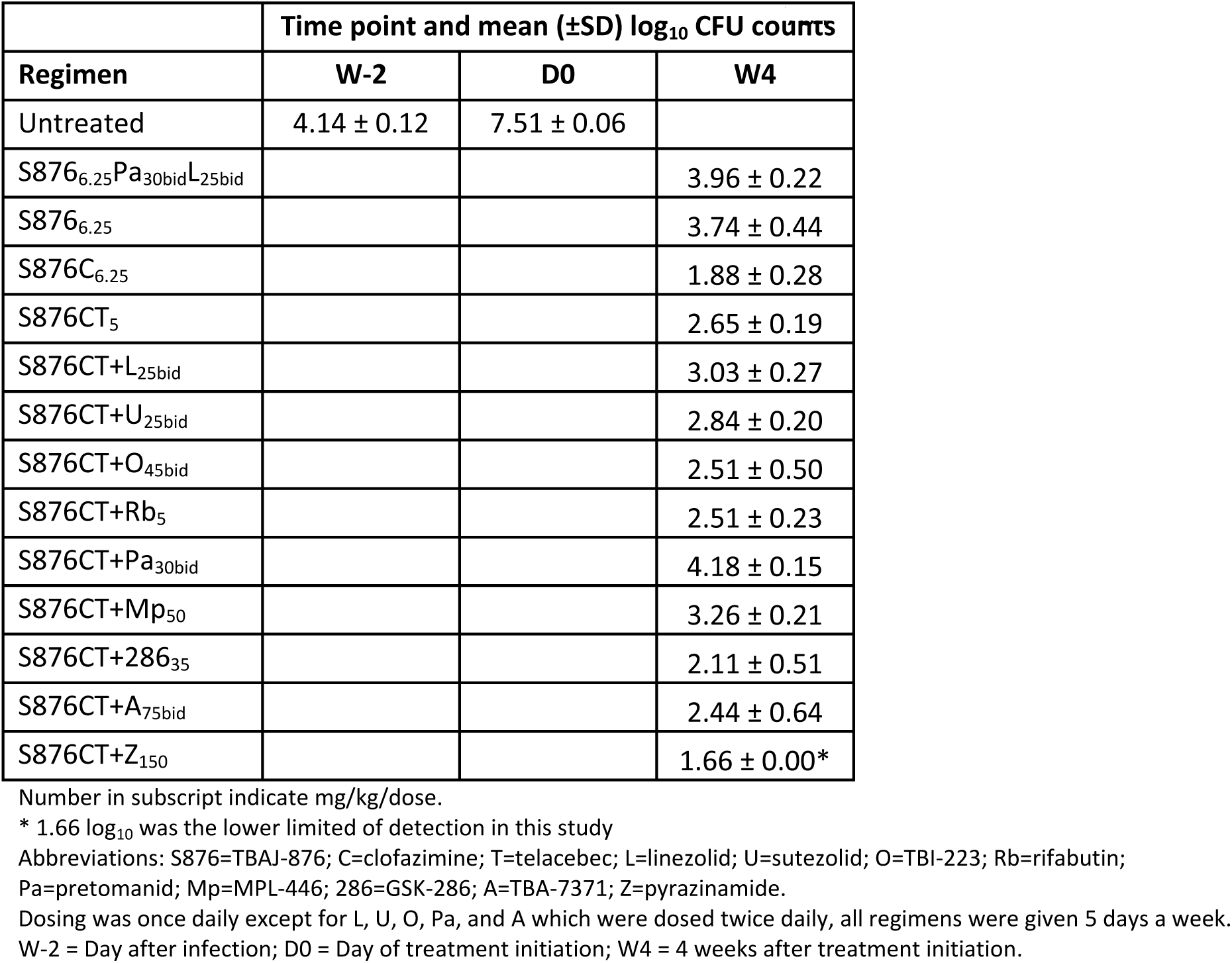
Mean CFU counts at week 4 in Experiment 2 (H37Rv)

### Experiment 3: Deconvoluting the effects of telacebec in combination with TBAJ-587, clofazimine and/or pyrazinamide

Combinations of BDQ with either CFZ or PZA were previously shown to have strong additive bactericidal and sterilizing activity in mouse models of TB (54-57). Experiment 3 was designed to evaluate the components of the DARQ-CFZ-T regimen tested in Experiment 2 and the interaction between T and the other next-generation DARQ in clinical development, TBAJ-587, with or without CFZ or PZA. To attempt to realize the synergy observed in vitro and gather information on the maximum possible effect, higher doses were tested for CFZ and T. A second objective was to determine if T would be as effective as TBAJ-587 when combined with either CFZ or PZA. The experimental scheme is shown in **Table S3**. BALB/c mice received a high-dose aerosol infection with *M. tuberculosis* H37Rv and were initiated on treatment 2 weeks post-infection. Monotherapy arms were limited to 4 weeks of treatment due to risk of resistance selection with longer treatment. Arms containing TBAJ-587 and PZA were limited to 4 weeks of treatment because CFU counts were expected to be very low or undetectable at 8 weeks and unlikely to permit assessment of the contribution of T. Other arms were evaluated after 4 and 8 weeks of treatment.

*The mean CFU count in the lungs at D0 was 7.52 ± 0.13 (*Table 2. Mean CFU counts at week 4 in Experiment 2 (H37Rv)

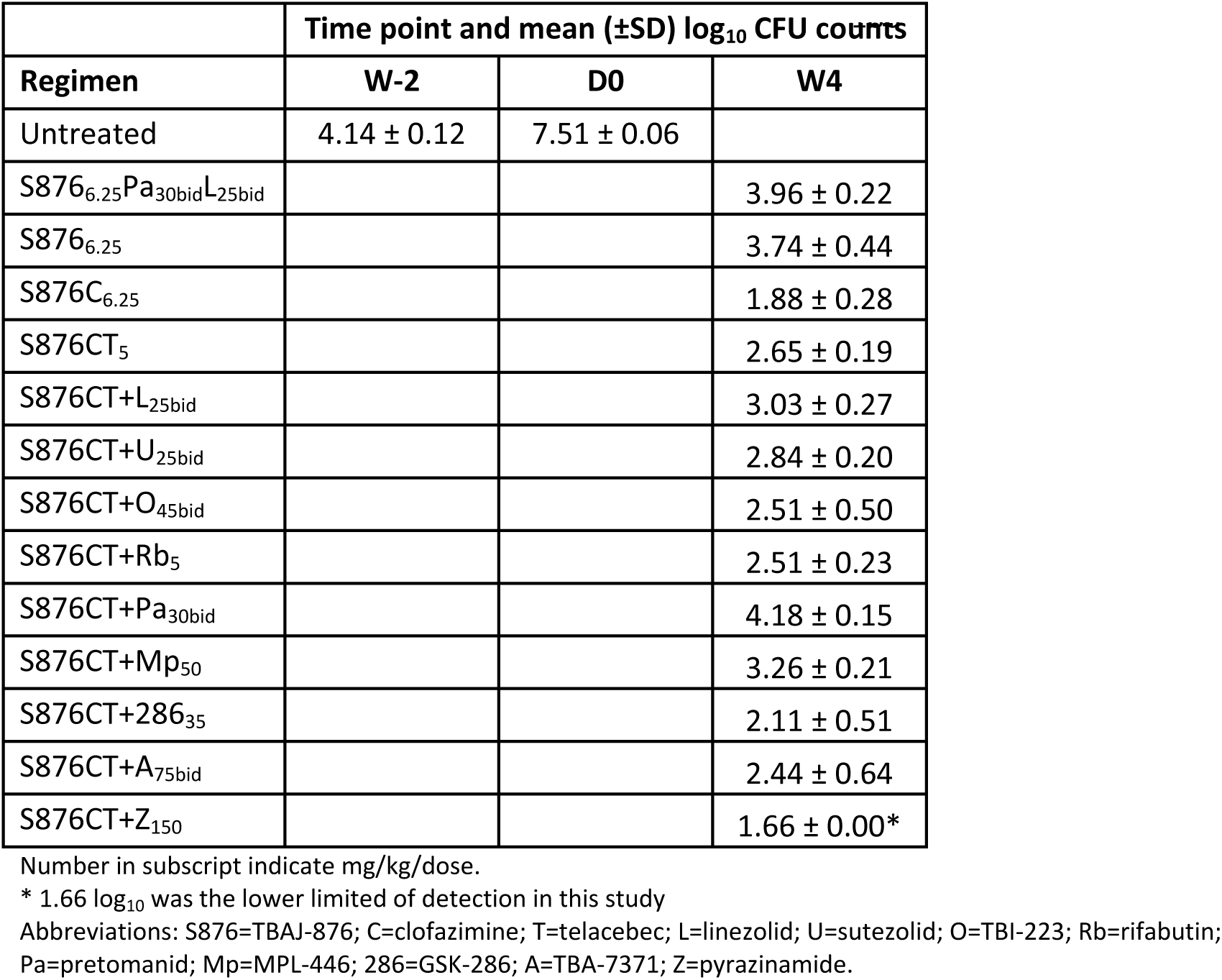
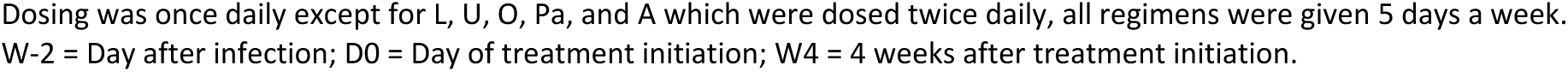

**Table 33**). Untreated mice experienced an increasing bacterial burden and met humane endpoints for euthanasia at Week 3. Single drug treatment with T at 10 mg/kg and 50 mg/kg was bacteriostatic, preventing death in T-treated mice, but not reducing lung CFU counts compared to D0 CFU counts. Furthermore, no dose-response was observed. TBAJ-587 alone at 12.5 mg/kg reduced the bacterial burden by more than 4 log_10_ over 4 weeks, comparable to the observed CFU reduction of TBAJ-876 monotherapy at 6.25 mg/kg in Experiment 2, and in line with a previous experiment in which TBAJ-587 at 25 mg/kg reduced the lung CFU count by approximately 5 log_10_ over 4 weeks (32). Addition of either PZA (p<0.001) or CFZ (p<0.01) significantly increased the bactericidal activity of TBAJ-587, resulting in the two most active drug combinations tested. In contrast, addition of T antagonized the activity of TBAJ-587 over the course of 4 weeks (p<0.01). No difference in the overall effect was observed for either dose of T in combination with TBAJ-587 after 4 or 8 weeks of treatment. Similarly, the addition of T to combinations of TBAJ-587 plus either CFZ or PZA also resulted in higher CFU counts after 4 and 8 weeks of treatment, although the effect of T on these combinations was not statistically significant. Whereas all mice treated with TBAJ-587 plus CFZ had no detectable CFU after 8 weeks of treatment, all mice treated with the same combination plus T remained culture-positive at this time point. Replacing TBAJ-587 with T in combinations containing CFZ or PZA resulted in significantly higher CFU counts. Compared with T monotherapy, the addition of PZA reduced the mean CFU count by over 3 log_10_ and the addition of CFZ reduced the mean CFU count by around 1 log_10_, but we did not assess whether these combinations with T were more effective than PZA or CFZ alone.

To determine if the observed antagonistic effect of T on DARQ-containing combinations could be due to drug-drug interactions which reduce drug exposures, steady state plasma concentrations of TBAJ-587 (**Table S4a**), its active M3 metabolite (M3) (**Table S4b**), and T (**Table S4c**) were measured to compare the exposures of the drugs alone and in combination with companion drugs. For TBAJ-587 and its M3 metabolite, the mean concentrations were compared between mice administered TBAJ-587 alone and mice administered TBAJ-587 in combination with T 10 mg/kg (T_10_), T 50 mg/kg (T_50_), and T_10_ plus CFZ. For T, the mean concentrations were compared between mice administered T_10_ alone and T_10_ in combination with TBAJ-587 and TBAJ-587+CFZ, as well as between mice administered T_50_ alone and T_50_ in combination with TBAJ-587.

Mean concentrations of TBAJ-587 and the M3 metabolite did not differ significantly between most groups and time points assessed. However, the mean concentrations of TBAJ-587 were significantly lower at 1h and 5h when TBAJ-587 was administered in combination with T_10_+C compared to treatment with TBAJ-587 alone (p<0.05). Similar results were observed for the M3 metabolite of TBAJ-587; at the 5h timepoint, the mean concentration in the TBAJ-587+T_10_+C group was significantly lower than that in the TBAJ-587 alone group (p<0.05). Furthermore, a significantly higher mean concentration of M3 was observed in the TBAJ-587+T_50_ group compared to TBAJ-587 alone (p<0.05) or TBAJ-587+T_10_ (p=0.001). A significantly lower mean concentration of T at 5h post-dose was observed in the group receiving TBAJ-587+T_10_ compared to the group receiving T_10_ alone (p<0.05). No other significant differences in T concentrations were noted between groups.

### Experiment 4: Evaluating the contribution of telacebec to combination drug regimens against *M. tuberculosis* HN878

Recent work suggests that the HN878 strain of *M. tuberculosis* is more susceptible to T than the H37Rv strain, likely because the latter more effectively utilizes the compensatory cytochrome bd oxidase to reduce telacebec’s effectiveness (49). To investigate if this differential vulnerability affects the contribution of T to novel drug combinations *in vivo*, we infected BALB/c mice with HN878 and evaluated T with a variety of companion drug combinations, including some of those previously tested against H37Rv in Experiment 1. Both BDQ-containing and BDQ-sparing regimens were included to investigate the interaction between BDQ and T against HN878. The scheme of the experiment is shown in **Table S5**.

The results confirmed the hypothesis that the HN878 strain is more susceptible to T-containing combinations than the H37Rv strain. Unlike the antagonism observed when T was added to TBAJ-876+CFZ and TBAJ-587+CFZ against H37Rv in Experiments 2 and 3, respectively, the addition of T to BDQ+CFZ significantly increased the bactericidal effect against HN878 in Experiment 4 (p<0.0001) **(Table 4)**. Unlike the antagonistic effect of adding T to PaL against H37Rv, the same addition against HN878 resulted in a statistically non-significant lower mean CFU count. Lastly, the addition of T to PMZ resulted in a statistically significant additive effect against HN878 (p<0.01) but not H37Rv. A significant additive effect of T with BPaC was also observed (p<0.0001), and the 4-drug BPaCT was much more effective than BPaL±T. No significant effects were observed with addition of T to the combinations of BPaL, BPaM, BGA and PaRb. The effect of adding T to the BCM combination could not be evaluated because all BCM-treated mice were culture-negative by the end of one month of treatment and the addition of T did not change this result. Adding T to BMZ increased the average CFU count significantly (p<0.05) and increased the percentage of mice with culturable bacteria from 25% to 75%.

**Table 3.**
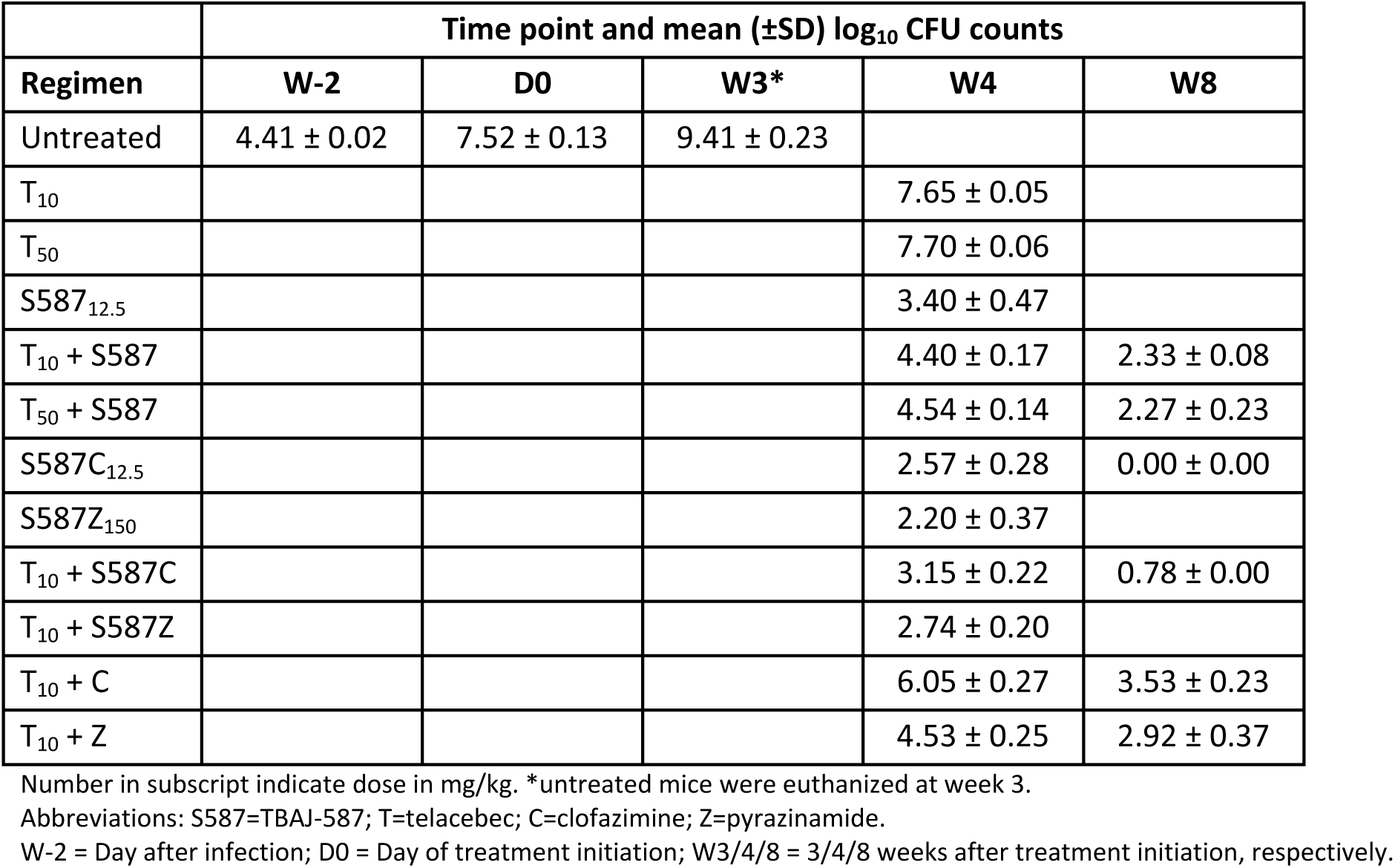
Mean CFU counts at week 4 and week 8 in Experiment 3 (H37Rv)

**Table 4.** Mean CFU counts at week 4 in Experiment 4 (HN878)

| | Time point and mean (±SD) $\log_{10}$ CFU counts | | |
| --- | --- | --- | --- |
| Regimen | W-2 | D0 | W4 |
| Untreated | 4.13 ± 0.04 | 6.73 ± 0.10 | |
| B<sub>25</sub>C<sub>6.25</sub> | | | 3.12 ± 0.36 |
| BCT<sub>5</sub> | | | 0.41 ± 0.70 |
| BCM<sub>100</sub> | | | 0.00 ± 0.00 |
| BCMT | | | 0.00 ± 0.00 |
| Pa<sub>50</sub>L<sub>100</sub> | | | 6.31 ± 0.23 |
| BPaL | | | 4.04 ± 0.21 |
| BPaLT | | | 3.43 ± 0.49 |
| TPaL | | | 6.05 ± 0.21 |
| BPaC | | | 2.50 ± 0.28 |
| BPaCT | | | 0.39 ± 0.45 |
| BPaM | | | 2.90 ± 0.12 |
| BPaMT | | | 3.17 ± 0.16 |
| BMZ<sub>150</sub> | | | 0.20 ± 0.39 |
| BMZT | | | 0.94 ± 0.66 |
| B286<sub>35</sub>A<sub>200</sub> | | | 0.00 ± 0.00 |
| B286AT | | | 0.57 ± 0.66 |
| PaRb<sub>5</sub> | | | 6.38 ± 0.32 |
| PaRbT | | | 6.00 ± 0.20 |
| P<sub>10</sub>MZ | | | 5.96 ± 0.23 |
| PMZT | | | 4.77 ± 0.12 |

Number in subscript indicate mg/kg/dose.
Abbreviations: B=bedaquiline; C=clofazimine; T=telacebec; M=moxifloxacin; Pa=pretomanid; L=linezolid; Z=pyrazinamide; 286=GSK'286; A=TBA-7371; Rb=rifabutin; P=rifapentine.
W-2 = Day after infection; D0 = Day of treatment initiation; W4 = 4 weeks after treatment initiation.

### Experiment 5: Determination of telacebec MICs against *M. tuberculosis* H37Rv strain and isogenic *Rv0678* mutants

Given the increasing recognition of *Rv0678* mutations associated with baseline and acquired phenotypic resistance to BDQ and CFZ (27, 28), and the impact of such mutations on a number of other TB drugs, including new DARQs (18-26), we investigated whether *Rv0678* mutations associated with BDQ resistance also affect the susceptibility of *M. tuberculosis* to T by determining the MIC of T against a pair of isogenic *Rv0678* mutants using the agar proportion method. The MIC of T was 8 ng/ml against the parental H37Rv strain and the Rv0678_2 mutant, which is isogenic except for a single nucleotide insertion Gly65 (+G) between nucleotides 193 and 194 of the *Rv0678* gene (18). Against the Rv0678_J4 mutant, which is also isogenic except for an IS6110 insertion between nucleotides 49 and 50 of the *Rv0678* gene (18), the T MIC was 4 ng/ml.

## Discussion

The current study used a well-established BALB/c mouse model of TB to evaluate the contribution of the first-in-class QcrB inhibitor T to the bactericidal activity of novel drug combinations targeting mycobacterial respiration and oxidative phosphorylation against two different strains of *M. tuberculosis*. Given the clinical success of the ATP synthase inhibitor BDQ and the ongoing clinical development of the more potent next-generation DARQs TBAJ-587 and TBAJ-876, we explored the effect of combining T with DARQ-containing combinations *in vivo*. Having previously observed additive bactericidal and sterilizing activity of BDQ and CFZ in this model and given the evidence of synergistic activity of BDQ, CFZ and T *in vitro* (48), we were particularly interested in the interactions of these 3 classes *in vivo*. To our knowledge, this is the first report describing the effects of this combination of drugs acting on the respiratory chain in a murine model of TB.

Against *M. tuberculosis* H37Rv (Experiments 2 and 3), TBAJ-876 and TBAJ-587 individually showed strong bactericidal activity, reducing lung CFU counts by around 4 log10 compared to D0 counts, results which are in line with previously published data (16, 27). The addition of CFZ increased the activity of either DARQ by approximately 1-2 log10, similar to previous observations with addition of CFZ to BDQ in this model (48). The observed bacteriostatic effect of T monotherapy was also consistent with previous *in vitro* results (42). Previous results in mouse infection models have varied, with some finding bactericidal activity (29) and others observing only bacteriostatic effects (49). Surprisingly, the addition of T antagonized or, at least, did not increase, the activity of either DARQ+CFZ combination, and also antagonized the activity of TBAJ-587 when TBAJ-587+T was compared to TBAJ-587 alone. Some *in vitro* studies have shown lack of additivity or antagonism when T or another QcrB inhibitor is added to BDQ (42, 50). However, the addition of T to a DARQ+CFZ combination was expected to be additive (42).

In stark contrast to the results observed against infections with the H37Rv strain, we found a strong additive effect when T was combined with a DARQ (BDQ) and CFZ against an infection with the HN878 strain. The addition of T to the combination of BDQ, Pa, and CFZ (BPaCT) was also advantageous against HN878 infection, reducing the mean CFU count by over 2 log_10_ compared to BPaC alone in Experiment 4. Whereas BPaCT reduced the mean CFU count by over 6 log_10_ compared to D0 and was superior to BPaL against HN878 in Experiment 4, TBAJ-876 plus PaCT reduced the mean CFU count by only 3.33 log_10_ and was significantly worse than TBAJ-876 plus PaL against H37Rv in Experiment 2. A similar strain-dependent differential contribution of T emerged when T was added to the PaL backbone, with T significantly antagonizing this DARQ-sparing backbone and increasing the mean CFU count by 1.05 log_10_ against H37Rv in Experiment 1, but lowering the mean CFU count, albeit non-significantly, against HN878 in Experiment 4.

BPaMZ was recently tested as a 4-month regimen for drug-sensitive TB in the SimpliciTB trial (ClinicalTrials.gov identifier NCT03338621), but did not meet non-inferiority criteria when compared to the 6-month standard-of-care due to treatment discontinuations from adverse events. Therefore, we tested whether T could replace Pa or Z, two components thought most to be associated with hepatoxicity in this combination (58). We observed similar results between the strains when T was used in place of Pa in BPaMZ. In both strains the addition of T was slightly antagonistic to the BMZ backbone. Against HN878 infection, addition of T to BPaM had no significant effect.

A regimen combining PMZ with isoniazid is thus far the only 4-month regimen proven to be non-inferior to the 6-month first-line regimen for drug-susceptible TB. However, the utility of isoniazid is limited by the prevalence of isoniazid monoresistance. As QcrB inhibitors have shown promising interactions with rifamycins and PZA (59), we compared the contribution of T to the PMZ backbone against both *M. tuberculosis* strains. Whereas the addition of T to PMZ had no significant effect during the first month against H37Rv, it reduced the mean CFU count by over 1 log_10_ against HN878. Additional studies using relapse as an endpoint and in a C3HeB/FeJ mouse model that develops caseating lung lesions may be useful to further explore the potential value of incorporating T into the BPaMZ and PMZ regimens.

The findings regarding *M. tuberculosis* strain-dependent contributions of T discussed above reinforce other recent findings from this BALB/c mouse model that demonstrated a differential contribution of linezolid to the BPa backbone against these two *M. tuberculosis* strains (60) and lead to one of our main conclusions—that important differences in drug susceptibility between different lineages and strains of *M. tuberculosis* exist and deserve greater attention in regimen development. With respect to the potential contribution of T to some of the novel regimens examined here, our observations require further investigation with additional strains from the two major lineages represented here by H37Rv and HN878 to determine the extent to which these are lineage-associated, versus strain-specific, effects and to better understand the mechanisms behind them. For example, we recently reported the considerable bactericidal and sterilizing activity of the B286A regimen in mice (13). The current results from Experiment 1 showing similar bactericidal activity of B286A and BPaL after 8 weeks of treatment in mice infected with H37Rv are in line with prior results. However, in Experiment 4 in which mice were infected with HN878, B286A rendered all mice culture-negative after 4 weeks of treatment and was clearly superior to BPaL (as well as PMZ) at this time point. We found that T did not add activity to the 286A backbone and therefore could not replace any portion of the BDQ contribution to B286A in Experiment 1, but we did not evaluate the contribution of T to 286A against HN878. Studies to further investigate the B286A backbone and the extent to which T can add activity or replace BDQ (with or without concomitant use of CFZ) should consider inclusion of additional *M. tuberculosis* strains from lineage 4 in particular.

The antagonistic effects of T on the activity of the DARQ-containing, and especially DARQ+CFZ-containing, regimens we observed in H37Rv-infected mice were surprising. Analysis of plasma drug concentrations in Experiment 3 did not provide compelling evidence of a drug-drug interaction between TBAJ-587 and T that would explain the antagonism, as the concentrations of TBAJ-587 and its active M3 metabolite were not consistently significantly lower in mice receiving TBAJ-587+T compared to those receiving TBAJ-587 alone. Only the M3 concentrations measured at 5 h post-dose in the TBAJ-587+T_50_ arm were significantly lower than the corresponding concentrations measured in the TBAJ-587 alone arm. However, TBAJ-587 and M3 concentrations were consistently significantly lower in mice receiving TBAJ-587+T_10_+CFZ compared to groups receiving TBAJ-587 alone or in combination with T, despite the fact that the addition of CFZ to this arm significantly increased its bactericidal activity over TBAJ-587+T_10_ only. Because drug concentrations were not assessed in the TBAJ-587+CFZ arm and CFZ concentrations were not measured in any arm, we were unable to determine if co-administration with CFZ itself lowered TBAJ-587 concentrations or if co-administration with both T and CFZ was necessary; nor could the impact of T on CFZ concentrations be assessed. The strong additive effects observed when T was added to BDQ+CFZ-containing regimens in HN878-infected mice provided additional, albeit indirect, evidence that the antagonism was not due solely to a drug-drug interaction that lowered DARQ concentrations.

Since a pharmacokinetic interaction did not appear to explain the antagonistic effect of T on DARQ activity against H37Rv, other possible explanations for this observation should be considered. An explanation may lie in the considerable respiratory flexibility of *M. tuberculosis* and the time course of its adaptation during mouse infection. *M. tuberculosis* is known to rely on the bd oxidase, an alternative non-proton-pumping terminal oxidase, to mitigate the effects of T-mediated blockade of the bc_1_:aa_3_ oxidase (61, 62), but upregulation of bd oxidase expression also enables an increase in ETC flux without contributing to membrane hyperpolarization in the face of blockade of ATP synthase by BDQ (61). Indeed, loss of the bd oxidase renders *M. tuberculosis* more susceptible to BDQ *in vitro* and *in vivo (61, 63)*. Temporal transcriptional profiling upon mouse infection has shown that *M. tuberculosis* also adapts to the onset of Th1 cell-mediated immunity *in vivo* by upregulating the bd oxidase and decreasing dependence on the bc_1_:aa_3_ oxidase (64). Unlike the standard *in vitro* conditions, this alteration would lead to lower expression of bc_1_:aa_3_ oxidase compared to bd oxidase prior to any treatment, which could reduce the benefit of T inhibition of the former while still promoting further additive expression of bd oxidase. Less consistent with prior *in vitro* results was our finding that addition of T to DARQ+CFZ combinations led to antagonism or, at least, lack of additive effects in mice infected with H37Rv. Indeed, Lamprecht et al. previously showed that addition of T to BDQ+CFZ increased bactericidal activity *in vitro* against H37Rv in axenic media and in infected RAW264.7 cells and proposed a mechanistic model whereby BDQ and T increase ETC flux via bd oxidase, which potentiates the bactericidal mechanism of CFZ to transfer electrons from NDH2 to oxygen to produce ROS (48). However, the additive effect of T *in vitro* in those experiments was relatively modest and the same imbalance of bd oxidase activity relative to bc_1_:aa_3_ oxidase activity *in vivo* as described above could mitigate against this small effect in mice. Since Shi et al. demonstrated that bd oxidase expression is highest during the onset of the adaptive immune response in mice and then returns to baseline as chronic infection is established (64), it may be worthwhile to revisit these combinations in a more chronic mouse infection model.

We hypothesized that a fourth drug inhibiting transcription or translation would limit the ability of *M. tuberculosis* to adapt and compensate for actions of the DARQ+CFZ+T combination and therefore increase the susceptibility to the regimen. Indeed, each of these classes works well together with rifamycins, in particular. This theory was not confirmed in our experiments, as neither mechanistic class of inhibitors was able to significantly increase the activity of the backbone regimen in Experiment 2, possibly due to the previously described over-expression of bd oxidase during infection occurring prior to drug exposure. Pa, as well as DprE1 and MmpL3 inhibitors have shown some ability to augment the activity of BDQ-containing regimens (50, 65) in previous experiments. DprE1 inhibitors have also shown additive activity with T (66) and with CFZ (67). However, we did not observe additive effects when Pa, TBA-7371 or the bactericidal and orally bioavailable MmpL3 inhibitor MPL-446 were added to TBAJ-876+CFZ+T in our study. The only drug that added significantly to this 3-drug backbone was PZA, consistent with our prior observations of the potent bactericidal and sterilizing efficacy of BDQ+CFZ+PZA in this model (57, 68).

Finally, we found that *Rv0678* mutations that reduce *M. tuberculosis* susceptibility to BDQ do not affect the MIC of T. This encouraging result suggests that T may, as another ATP synthesis inhibitor, provide an option to augment or replace BDQ in regimens against infections caused by *Rv0678* mutants.

This study has limitations. First, we evaluated a limited set of drug combinations to test specific hypotheses based on prior *in vitro* and *in vivo* studies. The potential contribution of T to other combinations of TB drugs should be examined. Second, the potential effect on the DARQ and CFZ exposures when T was added to either DARQ+CFZ combination was not assessed, which prevents us from determining whether an adverse PK interaction explains at least part of the antagonistic effect of T in the 3-drug combination. However, the magnitude of the antagonistic effect of T on the 3-drug combination was comparable to that of its effect on the TBAJ-587+T combination, where no significant PK interaction was observed. Therefore, it is unlikely that a PK interaction explains the entire antagonistic effect. Third, we did not evaluate the contribution of T to the sterilizing activity of these regimens by assessing relapse prevention as an endpoint. This could be examined in future studies, but we are skeptical that a significant beneficial effect on relapse would have been observed against the H37Rv strain had the regimens been assessed using this endpoint, where there was no additive bactericidal effect. Fourth, our studies relied on a single strain of each of two lineages of *M. tuberculosis* and the results may not be generalizable to other strains in these or other lineages. We note that the HN878 strain (of Lineage 2) appears more vulnerable to T treatment and *qcrB* silencing via CRISPR interference than the H37Rv strain (of Lineage 4) (49) and it is possible that the laboratory-adapted H37Rv strain is something of an outlier in terms of its limited vulnerability to QcrB inhibition. Therefore, we believe it would be worthwhile to evaluate the contribution of T to similar regimens against additional Lineage 2 and 4 isolates, and isolates from other lineages as well. Finally, we used a BALB/c mouse infection model that does not develop caseating lung lesions. Further studies are warranted to evaluate the potential contribution of T to TB therapy, including combinations with a DARQ and CFZ, in a model that forms such necrotic lesions, such as C3HeB/FeJ mice.

In conclusion, T exhibited additive activity with BDQ+CFZ-containing combinations and the PMZ combination in a bacterial strain-dependent manner. These and other regimens deserve further evaluation against additional *M. tuberculosis* strains, as well as in sterilizing effect studies and C3HeB/FeJ mice to further assess the potential contribution of T to novel drug regimens. However, it seems likely that T may only reach its full potential when a suitable inhibitor of the alternative bd oxidase is available

## MATERIALS AND METHODS

### MIC determination

*M. tuberculosis* H37Rv (American Type Culture Collection strain ATCC 27294) and isogenic *Rv0678* mutants previously selected in the H37Rv background were grown in 7H9 media with 10% OADC and 0.05% Tween 80. The Rv0678_J4 mutant has an IS6110 insertion at nucleotide 49. The Rv0678_2 mutant has a guanine insertion between nucleotides 193 and 194 of the *Rv0678* gene. Both mutants were isolated during a previous study in our lab (18).

The agar proportion method was used to determine the MIC. A stock solution of T was prepared in DMSO (Fisher Scientific), serially diluted in 2-fold steps, and added (0.1% vol/vol) to 7H11 agar supplemented with 10% (vol/vol) OADC enrichment, and 0.5% (vol/vol) glycerol to achieve all doubling T concentrations between 1 ng/ml and 64 ng/ml. All the strains were grown in 7H9 medium with (0.05%) Tween and (10%) OADC to an OD_600nm_ = 1. The inoculum for MIC testing was prepared by adjusting this culture to approximately 10^5^ CFU/ml by diluting it 1:100. Undiluted inoculum and inoculum diluted 1:100 were seeded onto T-containing plates, while serial 10-fold dilutions up to 1:10,000 were seeded onto T-free plates to determine the CFU count of the inoculum. The MIC was defined as the lowest T concentration that prevented the growth of at least 99% of CFU compared to plates without T after 18 days of incubation at 37°C.

### Bacterial strain

*M. tuberculosis* H37Rv (American Type Culture Collection strain ATCC 27294) and *M. tuberculosis* HN878 were mouse-passaged and frozen at -80°C in aliquots. For each mouse infection, an aliquot was thawed, grown in liquid culture medium (Middlebrook 7H9) and then used to aerosol infect mice.

### Infection model

All animal procedures adhered to national and international guidelines and were approved by the Johns Hopkins University Animal Care and Use Committee. For all experiments 6-week-old female BALB/c mice were aerosol-infected with a culture of *M. tuberculosis* during log phase growth with an optical density at 600 nm of approximately 0.8-1.0 (D-14). Treatment was initiated 2 weeks later (D0). On D-13 and D0 mice were sacrificed for lung CFU counts to determine the number of bacterial implanted and CFU counts at the start of treatment.

### Media

Bacteria for the aerosol infection were cultured in Middlebrook 7H9 broth supplemented with 10% (vol/vol) oleic acid-albumin-dextrose-catalase (OADC) enrichment, 0.5% (vol/vol) glycerol, and 0.1% (vol/vol) Tween 80. Lung homogenates as well as the cognate 10-fold dilutions were plated on selective 7H11 agar (7H11 agar containing 50 μg/ml carbenicillin, 10 μg/ml polymyxin B, 20 μg/ml trimethoprim, and 50 μg/ml cycloheximide), supplemented with 10% (vol/vol) OADC enrichment, 0.5% (vol/vol) glycerol and 0.4% activated charcoal to adsorb any drug carried over in the homogenates (55, 69). Difco Middlebrook 7H9 broth powder, Difco Mycobacteria 7H11 agar powder, and BBL Middlebrook OADC enrichment were obtained from Becton, Dickinson and Company. Glycerol and Tween 80 were obtained from Fisher Scientific, and activated charcoal was obtained from J. T. Baker. All selective drugs were obtained from Sigma-Aldrich/Millipore-Sigma.

### Antibiotic treatment

In Experiment 1 mice were randomized to one of 17 treatment groups (**Table S31**), in Experiment 2 mice were randomized to one of 13 treatment groups (**Table S2**), in Experiment 3 mice were randomized to one of 11 treatment groups (**Table S33**), and in Experiment 4 mice were randomized to one of 20 treatment groups (**Table S4**). CFZ, TBAJ-587, TBAJ-876 were formulated in 20% hydroxypropyl-β-cyclodextrin solution acidified with 1.5% 1N HCl. T was prepared in 20% (wt/wt) d-α tocopheryl polyethylene glycol 1000 (Sigma) succinate solution. Pa was prepared in the CM-2 formulation as previously described (70). Rb and PZA were prepared in deionized H_2_0. Oxazolidinones (L, U and O) were prepared in 0.5% methylcellulose. G was prepared in 1% methylcellulose. TBA-7371 was prepared in 0.5% methylcellulose plus 0.1% Tween 80. MPL-446 was prepared in 15% Solutol HS 15 in 50mM Na-phosphate buffer at pH 6.5. Drugs were administered by gavage, 5 days per week. In Experiments 1 and 4, Pa 50 mg/kg and L 100 mg/kg were given once daily. In Experiment 2, Pa 30 mg/kg and L 25 mg/kg were given together twice daily, 8 h apart, resulting in a total daily dose of 60 mg/kg of Pa and 50 mg/kg of L. In Experiment 1 and Experiment 3 selected combination regimens were given for up to 2 months (**Table S3** and **Table S2**), in Experiment 2 and Experiment 4 all treatments were given for 1 month (**Table S3. Experimental scheme Experiment 3 (H37Rv)**

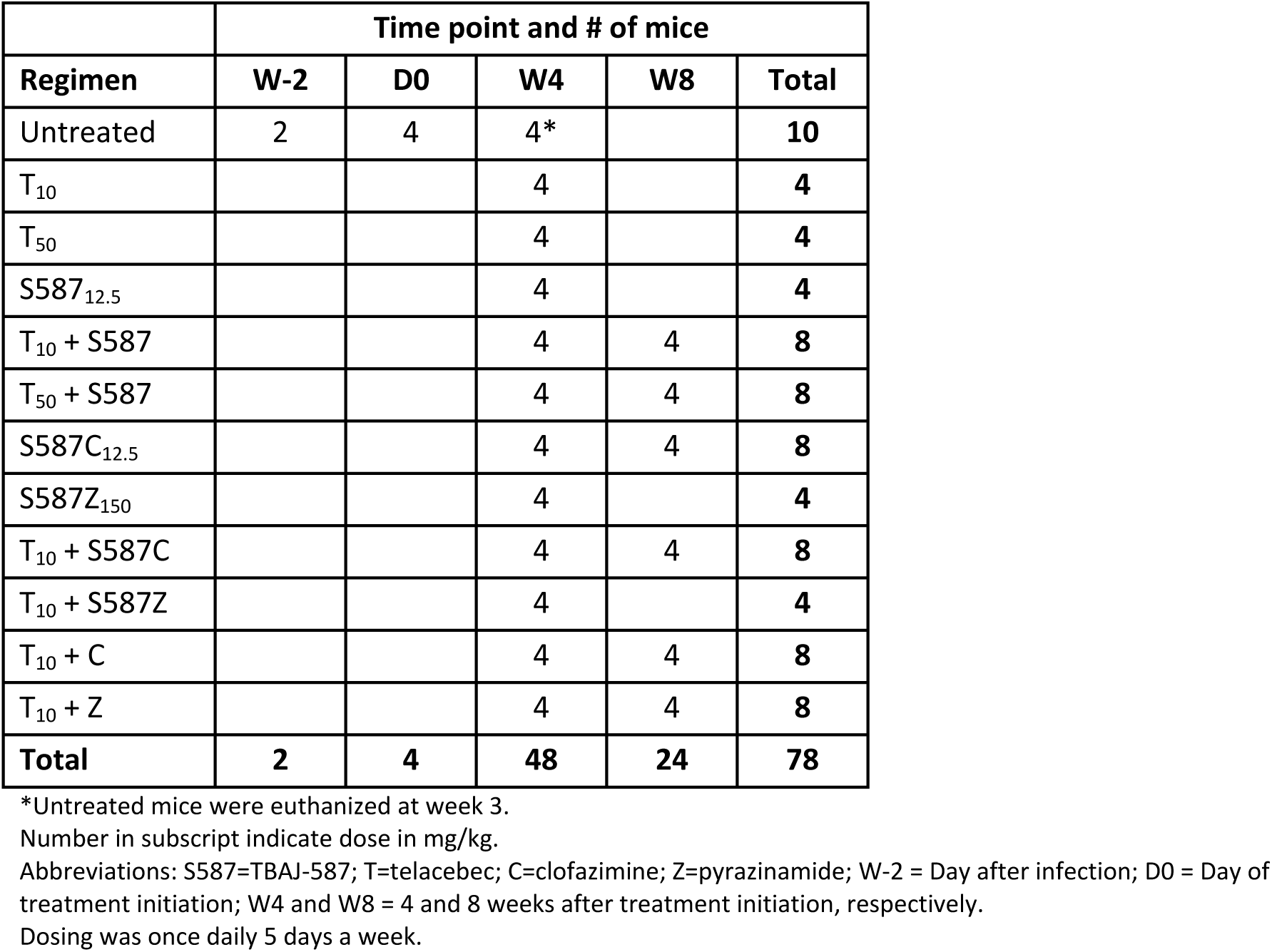

**3** and **Table S4**).

### Evaluation of drug efficacy in vivo

Efficacy was evaluated after 1 and 2 months of treatment by removing lungs aseptically and homogenizing in 2.5 ml PBS. Lung homogenates were then plated in serial dilutions on 7H11 agar plates supplemented with 0.4% charcoal and selective antibiotics (cycloheximide (20 μg/ml), carbenicillin (100 μg/ml), polymyxin B (400,000 U/ml), and trimethoprim (40 μg/ml)). CFU counts were performed after 4 and 6 weeks of incubation.

### Pharmacokinetics of TBAJ-587 and Telacebec

Multidose PK of T and TBAJ-587 in plasma was characterized in infected female BALB/c mice (Charles River Laboratories, Wilmington, MA) receiving oral doses once daily. In the fourth week of treatment in Experiment 3, 3 mice per group per time point were sampled by submandibular bleed at 1, 5, and 24 hours post-dose. Drug concentrations were quantified by a validated high-performance liquid chromatography/mass spectrometry method (Alliance Pharma Inc, Devault, PA).

### Statistical analysis

GraphPad Prism version 10.2 was used to compare group means by one-way ANOVA with Bonferroni’s correction to control for multiple comparisons.

## Acknowledgements

This research was supported by the TB Alliance.

## Tables

**Table S1.** Experimental scheme Experiment 1 (H37Rv)

|  | Time point and # of mice |  |  |  |  |
| --- | --- | --- | --- | --- | --- |
| Regimen | W-2 | D0 | W4 | W8 | Total |
| Untreated | 2 | 3 |  |  | 5 |
| Pa<sub>50</sub>L<sub>100</sub> |  |  |  | 4 | 4 |
| B<sub>25</sub>PaL |  |  |  | 4 | 4 |
| T<sub>5</sub>PaL |  |  |  | 4 | 4 |
| PaM<sub>100</sub>Z<sub>150</sub> |  |  | 4 |  | 4 |
| BPaMZ |  |  | 4 |  | 4 |
| TPaMZ |  |  | 4 |  | 4 |
| MZRb<sub>5</sub> |  |  | 4 |  | 4 |
| BMZRb |  |  | 4 |  | 4 |
| TMZRb |  |  | 4 |  | 4 |
| BMZ |  |  | 4 |  | 4 |
| BMZT |  |  | 4 |  | 4 |
| PMZ |  |  | 4 |  | 4 |
| PMZT |  |  | 4 |  | 4 |
| 286<sub>35</sub>A<sub>200</sub> |  |  |  | 4 | 4 |
| B286A |  |  |  | 4 | 4 |
| T286A |  |  |  | 4 | 4 |
| TC<sub>6.25</sub>286A |  |  |  | 4 | 4 |
| **Total** | **2** | **3** | **40** | **28** | **73** |

Number in subscript indicate dose in mg/kg. \*Untreated mice were euthanized at week 3.
Abbreviations: Pa=pretomanid; L=linezolid; B=bedaquiline; T=telacebec; M=moxifloxacin, Z=pyrazinamide; Rb=rifabutin; 286=GSK-286; A=TBA-7371; C=clofazimine. W-2 = Day after infection; D0 = Day of treatment initiation; W4 and W8 = 4 and 8 weeks after treatment initiation, respectively.
Dosing was once daily 5 days a week.

**Table S2.** Experimental scheme Experiment 2 (H37Rv)

|  | Time point and # of mice |  |  |  |
| --- | --- | --- | --- | --- |
| Regimen | W-2 | D0 | W4 | Total |
| Untreated | 2 | 3 |  | 5 |
| S876<sub>6.25</sub>Pa<sub>30bid</sub>L<sub>25bid</sub> |  |  | 5 | 5 |
| S876<sub>6.25</sub> |  |  | 5 | 5 |
| S876C<sub>6.25</sub> |  |  | 5 | 5 |
| S876CT<sub>5</sub> |  |  | 5 | 5 |
| S876CT+L<sub>25bid</sub> |  |  | 5 | 5 |
| S876CT+U<sub>25bid</sub> |  |  | 5 | 5 |
| S876CT+O<sub>45bid</sub> |  |  | 5 | 5 |
| S876CT+Rb<sub>5</sub> |  |  | 5 | 5 |
| S876CT+Pa<sub>30bid</sub> |  |  | 5 | 5 |
| S876CT+Mp<sub>50</sub> |  |  | 5 | 5 |
| S876CT+286<sub>35</sub> |  |  | 5 | 5 |
| S876CT+A<sub>75bid</sub> |  |  | 5 | 5 |
| S876CT+Z<sub>150</sub> |  |  | 5 | 5 |
| **Total** | **2** | **3** | **65** | **70** |

Number in subscript indicate mg/kg/dose.
Abbreviations: S876=TBAJ-876; C=Clofazimine; T=telacebec; L=Linezolid; U=Sutezolid; O=TBI-223; Rb=Rifabutin;
Pa=Pretomanid; Mp=MPL-446; 286=GSK-286; A=TBA-7371; Z=Pyrazinamide. W-2 = Day after infection; D0 = Day of treatment initiation; W4 = 4 weeks after treatment initiation.
Dosing was once daily except for L, U, O, Pa, and A which were dosed twice daily, all regimens were given 5 days a week.

**Table S3.**
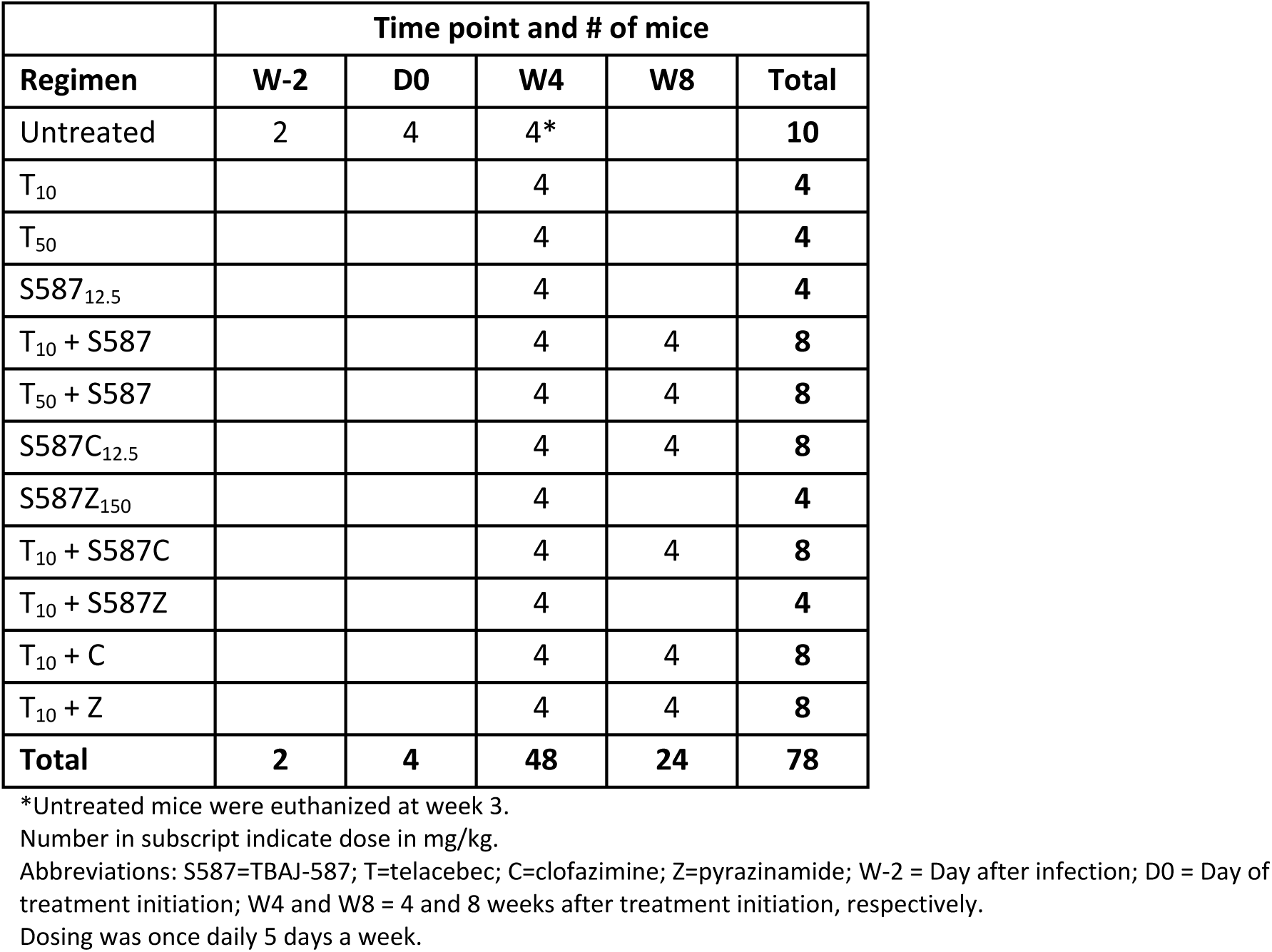
Experimental scheme Experiment 3 (H37Rv)

**Table S4a.** Mean concentrations of S587 at steady state.

|  |  | Regimen |  |  |  |
| --- | --- | --- | --- | --- | --- |
|  |  | S587<sub>12.5</sub> | S587+T<sub>10</sub> | S587+T<sub>50</sub> | S587+T<sub>10</sub>C<sub>12.5</sub> |
| Time-point | 1h | 794±323 | 465±115 | 331±373 | 93±57 |
|  | 5h | 411±151 | 242±146 | 424±216 | 120±62 |
|  | 24h | 310±158 | 264±165 | 279±206 | 75±16 |

Concentration of S587 in ng/ml ± SD. Number in subscript indicate mg/kg/dose.
Abbreviations: S587=TBAJ-587; T=telacebec; C=clofazimine

**Table S4b.**
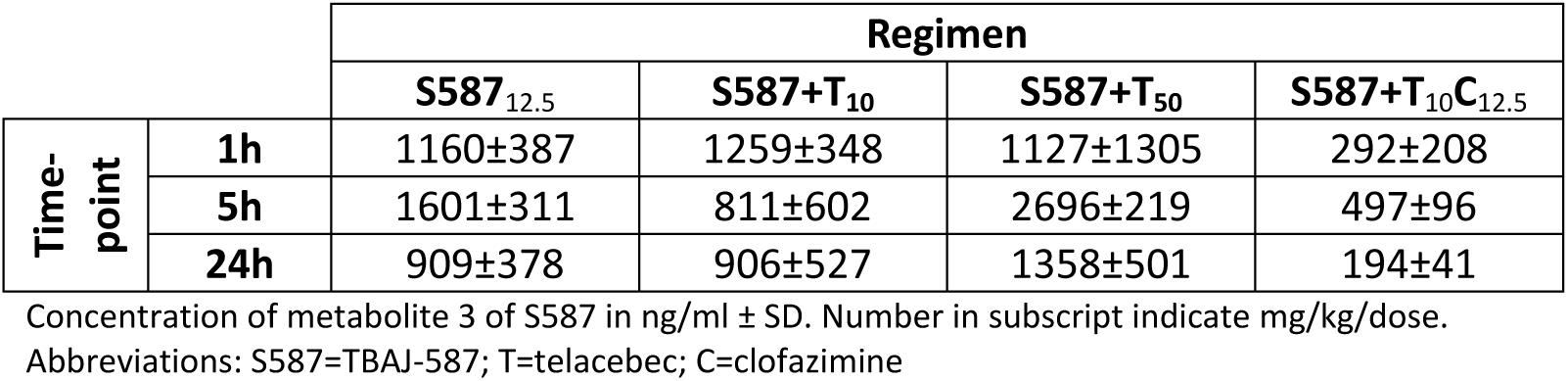
Mean concentrations of M3 metabolite of S587 at steady state.

|  |  | Regimen |  |  |  |
| --- | --- | --- | --- | --- | --- |
|  |  | S587<sub>12.5</sub> | S587+T<sub>10</sub> | S587+T<sub>50</sub> | S587+T<sub>10</sub>C<sub>12.5</sub> |
| Time-point | 1h | 1160±387 | 1259±348 | 1127±1305 | 292±208 |
|  | 5h | 1601±311 | 811±602 | 2696±219 | 497±96 |
|  | 24h | 909±378 | 906±527 | 1358±501 | 194±41 |

Concentration of metabolite 3 of S587 in ng/ml ± SD. Number in subscript indicate mg/kg/dose.
Abbreviations: S587=TBAJ-587; T=telacebec; C=clofazimine

**Table S4c.** Mean concentrations of T at steady state.

|  |  | Regimen |  |  |  |  |
| --- | --- | --- | --- | --- | --- | --- |
|  |  | T<sub>10</sub> | S587<sub>12.5</sub>+T<sub>10</sub> | S587+T<sub>10</sub>C<sub>12.5</sub> | T<sub>50</sub> | S587+T<sub>50</sub> |
| Time-point | 1h | 1462±435 | 1585±509 | 1445±1056 | 2642±1857 | 4608±6536 |
|  | 5h | 2265±102 | 861±644 | 1843±379 | 7310±3990 | 9881±2812 |
|  | 24h | 921±337 | 722±477 | 670±113 | 3008±330 | 2297±886 |

Concentration of Telacebec in ng/ml ± SD. Number in subscript indicate mg/kg/dose.
Abbreviations: S587=TBAJ-587; T=telacebec; C=clofazimine

**Table S5.** Experimental scheme for Experiment 4 (HN878)

| Regimen | Time point and # of mice |  |  |  |
| --- | --- | --- | --- | --- |
|  | W-2 | D0 | W4 | Total |
| Untreated | 2 | 3 |  | 5 |
| B<sub>25</sub>C<sub>6.25</sub> |  |  | 4 | 4 |
| BCT<sub>5</sub> |  |  | 4 | 4 |
| BCM<sub>100</sub> |  |  | 4 | 4 |
| BCMT |  |  | 4 | 4 |
| Pa<sub>50</sub>L<sub>100</sub> |  |  | 4 | 4 |
| BPaL |  |  | 4 | 4 |
| BPaLT |  |  | 4 | 4 |
| TPaL |  |  | 4 | 4 |
| BPaC |  |  | 4 | 4 |
| BPaCT |  |  | 4 | 4 |
| BPaM |  |  | 4 | 4 |
| BPaMT |  |  | 4 | 4 |
| BMZ<sub>150</sub> |  |  | 4 | 4 |
| BMZT |  |  | 4 | 4 |
| B286<sub>35</sub>A<sub>200</sub> |  |  | 4 | 4 |
| B286AT |  |  | 4 | 4 |
| PaRb<sub>5</sub> |  |  | 4 | 4 |
| PaRbT |  |  | 4 | 4 |
| P<sub>10</sub>MZ |  |  | 4 | 4 |
| PMZT |  |  | 4 | 4 |
| **Total** | **2** | **3** | **80** | **85** |

Number in subscript indicate mg/kg/dose.
Abbreviations: B=Bedaquiline; C=Clofazimine; T=telacebec; M=Moxifloxacin; Pa=Pretomanid; L=Linezolid; Z=Pyrazinamide; A=TBA-7371; Rb=Rifabutin. W-2 = Day after infection; D0 = Day of treatment initiation; W4 = 4 weeks after treatment initiation.
Dosing was once daily 5 days a week.

